# AF3 Inspector: In-Browser 3D Model Visualization and Confidence Analytics for AlphaFold 3

**DOI:** 10.64898/2026.09.21.753240

**Authors:** Xavier Gemis-Aebi, Franco Jiménez, Mirta G. Joveneche, Luciano A. Abriata

## Abstract

AlphaFold 3 (AF3) has broadened the computational structural biology landscape by predicting all-atom complexes across proteins, nucleic acids, small-molecule ligands, and ions, including chemical modifications. For non-experts, Google provides a fully web-based service to utilize AlphaFold 3. Unfortunately, despite a flexible and rich interface for inputs, the server’s outputs are incomplete and complicated for many users, as witnessed hands-on. Not all confidence metrics are displayed, and those that are shown do not present much interactivity; besides, only model number 1 is shown among the 5 produced by the software, and upon download the user finds the models are provided only in CIF format, which isn’t yet well-known especially among highly practical users. Here, we present the AlphaFold 3 Multi-Model Inspector (AF3 Inspector, https://pdbms.altervista.org/af3viewer/afviewer8.html), an open, client-side, zero-install web application designed to dissect, align, and interactively explore the complete ensemble of structural models produced by AlphaFold 3. Part of the PDB Manipulation Suite (https://pdbms.altervista.org/), the AF3 Inspector operates entirely within web browsers, meaning it’s available out of the box in all devices and operating systems. AF3 Inspector delivers 3D visualization in various styles and colors for all models, overlay with backbone alignment if requested, interactive plots for pLDDT profiles, pAE matrices and contact maps, chain-pair interaction heatmaps, automated detection of ligands, ions and post-translational modifications, and rapid mmCIF-to-PDB conversion allowing users to download the more familiar files. The platform addresses critical practical considerations highlighted in recent community benchmarks, such as CASP16, and extends the lightweight, client-side biophysical toolkit paradigm established by the PDB Manipulation Suite (PDBMS).

## Introduction

The transition from template-based homology modeling to deep learning has fundamentally reshaped structural biology.^1,2^ While AlphaFold 2 demonstrated that co-evolutionary embeddings from multiple sequence alignments could solve the majority of single-chain protein folds with experimental accuracy^3,4^, the release of AlphaFold 3 marked a departure from protein-only paradigms toward unified, all-atom biomolecular co-folding.^5^ By replacing the structural module with a generative diffusion network operating directly on raw atomic coordinates, AF3 routinely models multiprotein complexes alongside DNA, RNA, organic cofactors, therapeutic ligands, and post-translational modifications. Other contemporary deep learning systems subsequently adopted multimodal molecular inputs,^6–10^ and also built on top of the computer science unlocked by Deepmind’s AlphaFold 2 model to drive expansion toward proteome-scale structural genomics,^11^ structure analysis and searching,^12–14^ and ultimately protein design.^15–19^

However, in this new generative era of modeling, the fundamental bottleneck has shifted: generating coordinates is trivial, but critically evaluating structural plausibility, dynamic variability, and binding veracity is not if the right confidence scores are evaluated in the right way.^1^ When a run finishes, the AlphaFold server returns an archive containing five distinct structural models sampled by the diffusion process, accompanied by detailed JSON manifests of confidence scores. Structural biologists must immediately determine whether conformational differences across the five models capture genuine structural alternatives or stochastic noise in flexible loops, whether multi-chain assemblies form genuine physical interfaces or unproductively placed components, and how reliably non-protein entities—such as lipids or phosphorylated residues—have been docked. While overall fold topology is generally accurate across state-of-the-art predictors, multi-chain assemblies, ligand poses, and flexible regions demand deep scrutiny. In particular, global summary metrics like pTM and ipTM can mask severe localized packing failures or distorted binding pockets; then, dissecting residue-level pLDDT and pairwise pAE matrices is indispensable to separate trustworthy biological hypotheses from algorithmic hallucinations.

This gap between coordinate generation and practical interpretation is exacerbated by the current interface of the official AlphaFold Server. While the server (https://alphafoldserver.com/) provides an intuitive, feature-rich and highly flexible input builder capable of composing complex multi-chain assemblies with nucleic acids, organic ligands, ions, and diverse post-translational modifications, its output environment remains surprisingly constrained. In its current implementation, the web interface only displays a 3D visualization for the top-ranked model (Model 1, dubbed model 0 in the server’s output), leaving the remaining four structural hypotheses invisible unless the user downloads the full archive. Moreover, the graphical metrics presented on the server are largely static, offering minimal interactive cross-referencing between residue positions, PAE coordinates, and physical 3D contacts. Key diagnostic information such as the inter-chain minimum PAE matrix and token-level contact probabilities, are not exposed in an actionable manner. Compounding this challenge, the downloaded archives supply structural models exclusively in the newer mmCIF format. While mmCIF is the new established standard of the Worldwide Protein Data Bank, decades of legacy structural biology infrastructure, molecular dynamics packages, and practical bench workflows remain anchored to the classic PDB format. Moreover, for many wet-lab biologists and translational researchers, the PDB format is still the standard, and what’s worse, navigating command-line format conversions or inspecting complex JSON manifests simply to verify ligand binding interfaces or compare five alternate conformers creates a huge unnecessary barrier to adoption.

To address these analytical demands without forcing researchers through command-line scripts, Python environments, or third-party software to plot confidence metrics, visualize structural models and perhaps convert them to the more familiar PDB format, we have developed the AF3 Inspector, adopting the client-side architectural paradigm introduced in our PDB Manipulation Suite (PDBMS)^20^ and that we advocate^21–23^ for as materialized in most of our tools.^20,24–28^ The application, freely available without registration at https://pdbms.altervista.org/af3viewer/afviewer8.html, operates 100% within the user’s browser, handling archive extraction, coordinate parsing, alignment mathematics, format conversion, and matrix plotting entirely in local memory. This design guarantees multi-platform accessibility, zero software installation, and complete data confidentiality; in fact, the data never leaves the user’s machine.

### Architecture, workflow and implementation

The functional architecture of AF3 Inspector is organized into an end-to-end, browser-native analytical pipeline that operates without external server dependencies (Figure 1A). The workflow initiates when a user supplies an unmodified AlphaFold 3 output archive, either through a local file selector or via direct drag-and-drop onto the browser window. An internal decompression engine reads the archive in memory, indexing and extracting the job manifest (_job_request.json), per-model global confidence summaries (_summary_confidences_*.json), token-level interaction matrices (_full_data_*.json), and atomic coordinate files (_model_*.cif). Once parsed, the application populates two coordinated interface panels: a lateral control sidebar and a central analytical stage. The sidebar presents an executive summary of the prediction run, detailing entity stoichiometries, specified chemical modifications, and random seed parameters, followed by a comparative ranking table reporting pTM, ipTM, and overall ranking scores alongside direct, single-click PDB download links for each individual model. The central stage coordinates five interactive analytical tabs: an ensemble 3D molecular viewer driven by JSmol, a superposed multi-model pLDDT profile plotted across all five structures, interactive 2D heatmaps for token-level Predicted Aligned Error (PAE) and Predicted Contact Probabilities (< 8 Å), and an inter-chain interaction panel displaying minimum PAE and pairwise ipTM matrices for binder screening. Operating downstream of this visualization engine, a client-side conversion module translates mmCIF records into standardized PDB coordinates on the fly, feeding both the JSmol superposition pipeline and an automated archive generator that packages the converted PDB models, publication-grade PNG graphics, and an executive Markdown summary into a single downloadable ZIP archive.

**Figure 1.**
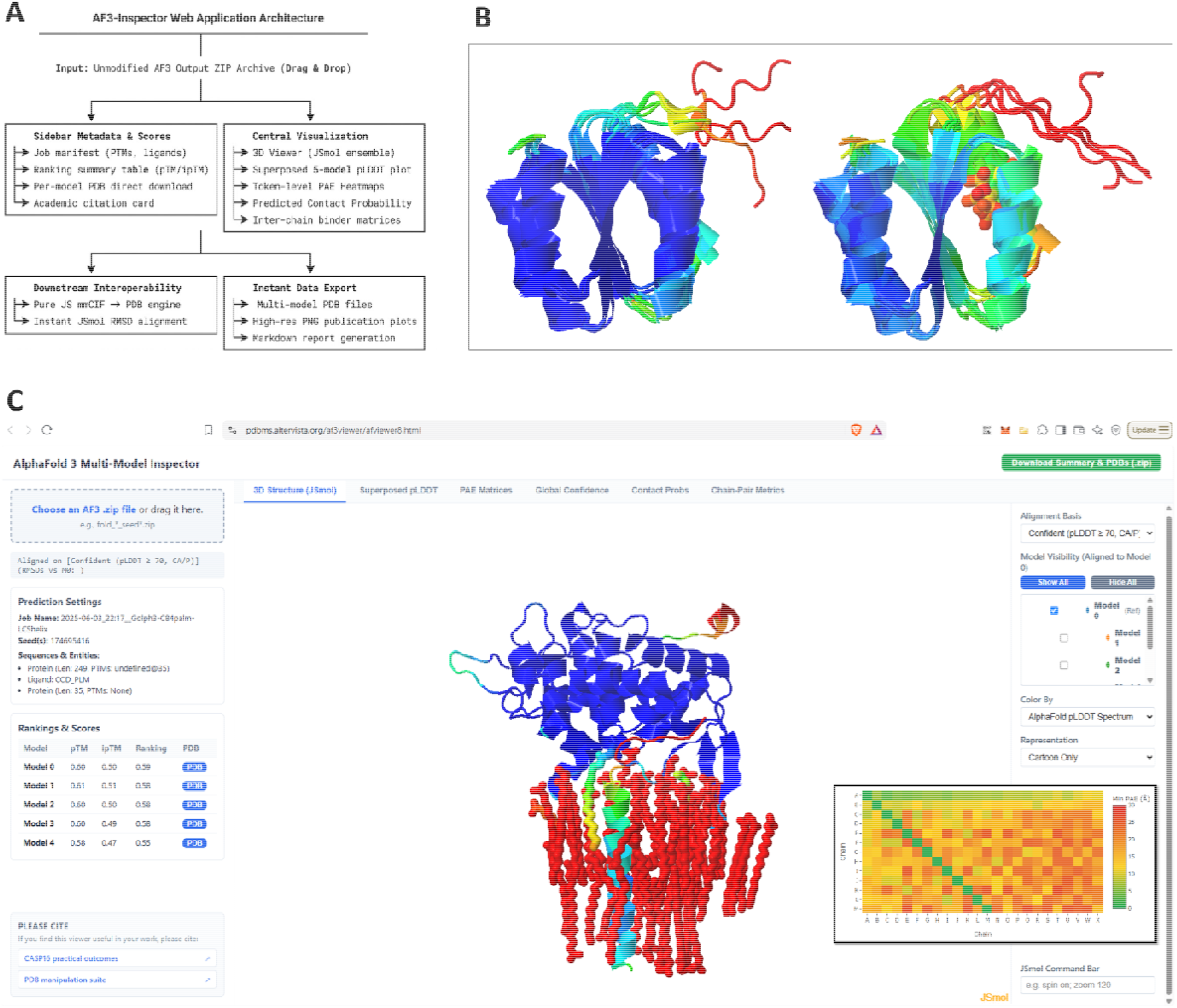
Operational workflow and representative structural biology applications of AF3 Inspector. **(A)** Overview of the client-side architecture. The user loads or drops a compressed AlphaFold 3 output ZIP file into the browser. The internal engine processes job metadata, token-level confidence arrays, and mmCIF models entirely in local memory, rendering synchronized multi-model JSmol viewports, overlaid 1D pLDDT traces, token-resolution PAE and contact probability heatmaps, and various matrices, all happening within the browser without server-side dependence (the website is only used to store the program, which runs entirely in the user’s device). **(B)** Application to understanding engineered dynamic switches as modeled by AF3: five models predicted for the protein in the non-phosphorylated (left) and phosphorylated (right) states, all colored by pLDDT after having aligned each to the first model. Notice how despite the high structural similarity, the models for the phosphorylated state have lowered confidence at and around the phosphorylated serine (shown as thick sticks). While this in principle means low confidence, it appears here to correlate with the regions most affected by phosphorylation as studied by NMR in the paper reporting this work. **(C)** The full AF3 Inspector web app, shown here with the user inspecting the top model for the complex predicted after feeding AF3 with two proteins (Golph3 with a palmitoylated cysteine plus a cognate peptide part of a transmembrane helix and 50 lipid molecules). The view recapitulates binding of GOLPH3 to the cognate peptide at the membrane interface, both proteins being modeled with intermediate to good confidence, the lipids ranked badly despite visually being correct in their placement, and with the palmitoylated cysteine showing high confidence near the Golph3 protein but lower confidence as it penetrates the membrane. In this panel, the inset is a small screenshot from the Chain-Pair Metrics tab, showing the chain-pair minimal pAE between pairs of chains; notice how the pAE is rather good for chains A-B (Golph3 with cognate peptide) and for chain A with many but not all lipids (chains C and beyond, likely those in contact with the protein).

### Structure-aware alignment for easier visual comparisons

Comparing five structural models sequentially without a unified spatial frame is disorienting because raw diffusion outputs have arbitrary rotational and translational orientations. Furthermore, naive all-atom superposition is routinely corrupted by flexible terminal tails or disordered loops. To resolve this, AF3 Inspector loads all five models as a synchronized multi-model ensemble in JSmol,^29^ used as the embedded web applet for molecular display, and applies JSmol’s internal quaternion *compare*() routine. The tool parses the B-factor column of Reference Model 0, isolates confident structural elements (defaulting to pLDDT ≥ 70 on Cα/P (protein/nucleic acid) backbone atoms), groups them into continuous sequence segments, and maps those identical residue ranges across all models. Disordered linkers and fluctuating termini are excluded from the mathematical superposition, yielding tight, domain-centric fits and reporting instantaneous RMSD values in the interface. Checkbox controls allow users to toggle any combination of models on and off without camera jumps, while a specialized color palette assigns coordinated colors to distinguish model divergence.

Free small-molecule ligands, prosthetic groups, and ions are selected through non-polymer queries and displayed as sticks and spheres. In parallel, by cross-referencing the parsed job specification and inspecting non-canonical residues in the coordinate stream, covalently attached post-translational modifications (PTMs) have their sidechains projected as sticks directly from the cartoon backbone.

### Multi-resolution confidence profiling, and format conversion

AF3 Inspector systematically renders the complete confidence hierarchy produced by AlphaFold 3. All five models have their residue-level pLDDT curves superposed on a single interactive canvas powered by Plotly.js, allowing researchers to more easily explore whether specific structural features and similarities or divergences across models coincide with local drops in model confidence. For pairwise interaction analysis, the app renders 2D heatmaps for both predicted Aligned Error (pAE, from 0 to 30 Å) and Contact probabilities (assuming contacts to hold when interatomic distances are under 8 Å). Accompanying explanatory banners clarify token resolution: while protein and nucleic acid chains are evaluated at residue centers (CA for protein residues, C1 for nucleic acid bases), small-molecule ligands and chemical modifications are tokenized on individual heavy atoms, yielding true single-atom resolution. For multimers, dedicated heatmaps display the minimum inter-chain PAE and pairwise ipTM matrices to evaluate binding interfaces. Finally, an embedded conversion engine transforms the mmCIF coordinates^30^ into standard PDB files, offering immediate per-model downloads as well as full archive export containing converted PDBs, publication-grade PNG plots, and markdown summaries.

### Practical Applications

To demonstrate the versatility of AF3 Inspector across distinct biophysical regimes, we examined two challenging modeling problems representing synthetic protein design and membrane-associated multiprotein signaling (Figure 1B-C). Following the operational workflow depicted in Figure 1A, these case studies highlight how simultaneous multi-model alignment and granular confidence dissection resolve critical questions.

The first application (Figure 1B) examines the de novo design of phosphorylation-induced protein switches recently reported by Buckley et al,^31^ where phosphorylation of a target serine is intended to drive a discrete conformational reorganization between active and inactive states. With a complex design strategy, Buckley et al managed to create a protein that exchanges between a closed state where a phosphosite serine is occluded and another state where it is exposed for phosphorylation, all validated by NMR spectroscopy and various other assays. Moreover, the work found that upon phosphorylation the serine-exposed state is much more exposed and the protein gains fast dynamics at the expense of losing slow, coordinated motions.

It turns out that when AF3 is asked to model the non-phosphorylated and phosphorylated versions of this protein, both tasks return 5 models where the protein is in the closed state with the serine/phosphoserine buried inside the core. However, deep inspection of the models as made easy by AF3 Inspector (in particular, here by loading each zip file on a separate browser tab) shows clearly how the pLDDT and other metrics fall largely at and around the target serine in the phosphorylated state (Figure 1B, right side) compared to the non-phosphorylated state.

The second application (Figure 1C) focuses on the peripheral membrane-targeting complex formed by Golgi phosphoprotein 3 (GOLPH3) bound to a cognate peptide that comes from the transmembrane helix of another protein, all in association with lipids as this happens at the membrane. The example was drawn from the molecular dissection by Theodoropoulou et al,^32^ based on modeling plus simulations and large experimentation.

Modeling peripheral membrane effectors with AlphaFold 3 presents a dual challenge: the algorithm must correctly configure multi-chain protein assemblies while simultaneously predicting the binding pose of non-polymer lipids and accounting for post-translational modifications like S-acylation. Within AF3 Inspector, the non-protein lipid headgroup is automatically identified and rendered in prominent stick-and-sphere representation nestled within the GOLPH3 binding pocket. The user can immediately cross-validate this visual binding pose against the inter-chain minimum PAE matrix (shown as an inset in Figure 1C towards the bottom right of the panel). The low chain-pair minimum PAE values between Golph3 and the cognate peptide and immediate lipids quantitatively verify a stable, high-confidence interface rather than a non-specific docking artifact, despite the bad pLDDT at some regions of the model.

## Conclusion

By unifying visualization with multi-resolution confidence metrics inside a very easy to use web application, AF3 Inspector should help bench biologists and modelers alike to more rapidly assess their models, especially when screening interfaces, evaluating ligand poses, exploring the effect of post-translational modifications, and generating research-ready PDB structures and figures in minutes.

## Code availability

The code is trivially accessible as the source code of the HTML page (opened with Ctrl+U in most browsers).

